# VCP inhibition prevents cone photoreceptor degeneration in the *cpfl1* mouse model of achromatopsia

**DOI:** 10.64898/2026.09.21.753131

**Authors:** Ana-Cristina Almansa-García, Bowen Cao, Anne-Sophie Petremann-Dumé, Ellen Kilger, Sylvie Bolz, Marius Ueffing, Blanca Arango-Gonzalez

## Abstract

Achromatopsia (ACHM) is a rare autosomal recessive retinal disorder characterized by absent cone photoreceptor function from early life, leading to severe visual impairment. Mutations in genes involved in the cone phototransduction cascade frequently result in elevated cyclic guanosine monophosphate (cGMP) levels and activation of stress pathways, including endoplasmic reticulum (ER) stress and the unfolded protein response. Targeting common downstream mechanisms rather than individual mutations may provide a broadly applicable therapeutic strategy. Here, we investigated whether pharmacological inhibition of valosin-containing protein (VCP), a key regulator of ER and protein homeostasis, can prevent cone degeneration in the spontaneous cone photoreceptor function loss 1 (*cpfl1*) mouse model of ACHM.

Organotypic culture of retinal explants from *cpfl1* mice were treated with the selective VCP inhibitor ML240. Cone survival, cell death, opsin expression and localization were assessed by TUNEL assay, immunohistochemistry, and quantitative image analysis.

ML240 treatment significantly increased cone density and improved cone opsin expression and trafficking to the outer segments (OSs) in *cpfl1* explants compared to controls. Importantly, rhodopsin trafficking in rod photoreceptors was unaffected, indicating that VCP inhibition did not impair normal rod phototransduction.

These findings demonstrate that VCP inhibition by ML240 effectively preserves cone photoreceptors and improves cone-specific functional markers in the *cpfl1* model. Targeting VCP may represent a mutation-independent therapeutic strategy for preventing cone death in ACHM.

## Introduction

Achromatopsia (ACHM) is a rare autosomal recessive retinal disease characterized by congenital or early-onset absence of cone photoreceptor function. Patients typically present with severely reduced visual acuity, photophobia, nystagmus, and impaired color discrimination [1]. Although its prevalence is low, ACHM causes lifelong visual disability due to its early onset and profound functional impairment [2].

Most of ACHM cases are caused by mutations in genes encoding components of the cone phototransduction cascade, including CNGA3, CNGB3, GNAT2, PDE6H, and PDE6C. In addition, mutations in ATF6, a regulator of endoplasmic reticulum homeostasis, have also been associated with the disease [2, 3]. Disruption of cone phototransduction frequently leads to sustained elevation of cyclic guanosine monophosphate levels [4-6], which has been associated with activation of multiple stress-related pathways and progressive cone degeneration. Although the precise mechanisms linking cGMP dysregulation to cone death remain incompletely defined, evidence from several models implicates proteostasis imbalance and stress-response signaling pathways as contributing factors [7].

Mutation-specific gene supplementation strategies have demonstrated promising safety profiles and variable functional improvement in preclinical models and early-phase clinical trials for CNGA3- and CNGB3-associated achromatopsia [8, 12]. Nevertheless, these approaches are mutation-specific and depend on the presence of viable cones, thereby limiting their clinical implementation and therapeutic window. An alternative strategy is to target common downstream pathways that mediate cone degeneration independently of the causative mutation.

Valosin-containing protein (VCP) is a multifunctional AAA+ ATPase that plays a central role in proteostasis, including ER-associated degradation, autophagy, and regulation of stress signaling pathways [13, 14]. Dysregulation of VCP-dependent processes has been implicated in multiple neurodegenerative conditions. Pharmacological inhibition and genetic modulation of VCP has previously been shown to attenuate photoreceptor cell death in models of inherited retinal degeneration, suggesting that modulation of proteostasis pathways may confer neuroprotection [15-19].

The spontaneous *cone photoreceptor function loss 1* (*cpfl1*) mutant mouse model carries a 16-bp insertion between exon 4 and 5 of the *Pde6c* gene, which codes for the cone photoreceptor cGMP phosphodiesterase catalytic alpha subunit, impairing the phototransduction cascade in cone photoreceptors [20]. This mouse model exhibits a disease phenotype similar to that observed in human patients with complete ACHM, including elevated cGMP levels, early-onset cone dysfunction, and progressive cone degeneration [6, 20].

The present study aims to evaluate the effect of the selective VCP inhibitor ML240 [21] on *cpfl1* mouse retinae, using an ex vivo organotypic retinal culture system. To determine whether VCP modulation can protect cone photoreceptors from degeneration, we assessed cone survival, structural integrity, opsin expression and localization.

## Methods

### Animals

Homozygous *cpfl1* mice of the line B6.CXBI-Pde6c^cpfl1^/J were kindly provided by Prof. M. Seeliger (Institute for Ophthalmic Research, Tübingen). C57BL/6J (WT; RRID:IMSR_JAX:000664) were purchased from The Jackson Laboratory and maintained in Tübingen. All animals were housed under standard white cyclic lighting conditions, with food and water provided ad libitum. Both male and female animals were included in the study.

All experimental procedures were in full compliance with the German Animal Welfare Act (Tierschutzgesetz) and approved by the local ethics committee (03/24 M). Special attention was paid to minimizing the number and suffering of animals.

### Retinal organ culture and treatments

*Cpfl1* or C57BL/6J mice eyes were enucleated at postnatal day 14 (PN14) under aseptic conditions and pretreated with 12 % proteinase K (0219350490, MP Biomedicals) for 15 minutes at 37 °C in R16 serum-free culture medium (07490743A, Invitrogen Life Technologies). The enzymatic digestion was stopped by the addition of 20 % fetal bovine serum (FBS; F7524, Sigma-Aldrich). The retina with attached RPE was dissected, and four radial cuts were made to flatten the tissue. Tissues were transferred onto a 0.4 μm polycarbonate membrane (Corning Life Sciences, CLS3412), with the RPE side facing the membrane. Cultured explants were maintained for 10 days in serum-free medium (DMEM/F12:DMEM, 1.5:1) supplemented with 2 % B27, 1 % N2, and 1 % antibiotic–antimycotic (Gibco). The medium was changed every two days. Treatment groups included control medium without any addition, 0.4 % DMSO (vehicle control, A994.1, Roth), and 20 µM ML240 (5153, Tocris).

### Histology

Retinal explants were fixed in a 4% paraformaldehyde (PFA) in 0.1 M phosphate buffer (PB, pH 7.4) for 45 minutes and cryoprotected in a sucrose gradient (10%, 20%, and 30%). Samples were embedded in cryomatrix (4583, Tissue-Tek® O.C.T. Compound, Sakura® Finetek, VWR) and snap-frozen in liquid nitrogen. Radial sections (14 µm) were cut, air-dried at 37ºC, and stored at -20ºC.

### TUNEL assay

Cell death was assessed using an in situ cell death detection kit (11684795910, Roche) conjugated with fluorescein isothiocyanate (FITC), according to the manufacturer’s protocol. Nuclei were counterstained with DAPI (1:5000 in PBS; D-9542, Sigma) for 5 minutes. Slides were mounted using Fluoromount-G (17984-25, Electron Microscopy Sciences).

### Immunohistochemistry

*Cpfl1* retina sections of 14 µm were washed three times for 5 minutes with PBS, permeabilized and blocked for 1 h at room temperature with 10% normal goat serum (311-053, PAA-Labs) or donkey serum (D9663, Sigma), plus 1% bovine serum albumin (K41-001, PAA-Labs) in 0.1% PBST (PBS 0.1% Tween). Afterwards, they were incubated overnight at 4°C with the respective primary antibodies (anti-cone arrestin 1:350, AB15282, Millipore; anti-M opsin 1:200, AB5405, Sigma Aldrich; anti-S opsin 1:300, sc-14363, Santa Cruz; anti-Rhodopsin 1:350, MAB5316, Sigma Aldrich) diluted in blocking solution. Secondary antibodies (Alexa Fluor™568 dye-conjugated goat anti-mouse IgG, 1:500, A11031, ThermoFisher; Alexa Fluor™ 568 dye-conjugated goat anti-rabbit IgG, 1:500, A11036, Molecular Probes) were applied for 1 h at room temperature, followed by DAPI counterstaining and mounting with Fluoromount-G (17984-25, Electron Microscopy Sciences).

### Microscopy and image analysis

Z-stack images were acquired using a Zeiss Axio Imager Z1 ApoTome microscope (20x or 63x objective) and analysed with Zen Blue 3.7. Three to four sections per sample were imaged. ONL rows were manually counted from DAPI staining. TUNEL-positive cells were quantified as a percentage of total ONL cells. The total ONL cell number was calculated by dividing the ONL area by the average photoreceptor nucleus size, which was obtained by measuring the area of at least six photoreceptor nuclei from different regions of the retinal sections using the contouring tool in Zen Blue 3.7. Cone density was calculated as cones per 100 µm of retinal section, averaging three different regions per image. Cone migration through the ONL was assessed by measuring the distance between the cone nucleus and the outer plexiform layer, for at least 100 cones per sample, and normalising for the thickness of the ONL. Rhodopsin distribution was measured by quantifying the ratio of rhodopsin mean fluorescence intensity between the ONL and the OS. All post-processing adjustments (contrast, brightness) were applied equally across groups.

### Statistical analyses

Analyses were performed using GraphPad Prism version 10. Data were presented as mean ± SEM. Each individual enclosed circle represents the average of an independent biological replicate (n). Group comparisons were performed using one-way analysis of variance (ANOVA), followed by Tukey’s multiple comparisons test. A p-value less than 0.05 was considered statistically significant.

## Results

### VCP inhibition by ML240 prevents primary cone degeneration in *cpfl1* mouse retinal explants

To determine the effect of VCP inhibition in the *cpfl1* mouse model, we employed an *ex vivo* organotypic culture system to culture mouse retinal explants as described previously [22, 23]. In short, eyes of *cpfl1* mice and the mouse line C57BL/6J, used as a WT control, were enucleated at postnatal day 14 (PN14), at the beginning of retinal degeneration in *cpfl1* mice [6]. Retinas were dissected with the retinal epithelium layer still attached and kept in culture for ten days *in vitro* until the peak of degeneration in *cpfl1* mice at PN24 [6]. To assess the effect of VCP inhibition, 20 µM of ML240 was administered in the culture medium. WT-type or mutant retinas, untreated or treated with the vehicle alone (0.4% DMSO), were used as controls (**Figure 1**).

**Figure 1.**
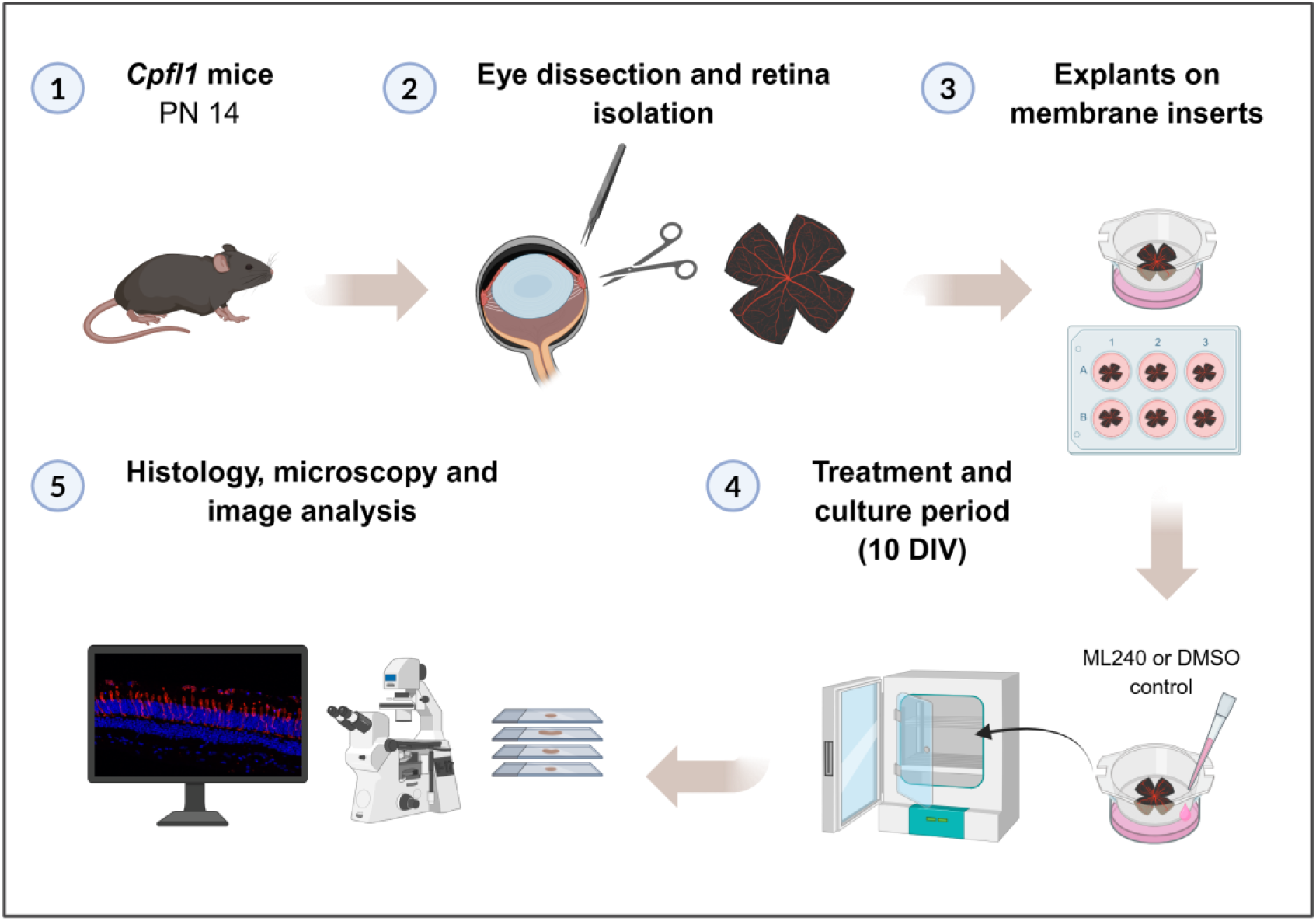
Schematic representation of the experimental procedure. (**1**) Eyes from *cpfl1* mice were enucleated at postnatal day 14 (PN14) and were dissected to isolate the neuroretina with the retinal pigment epithelium layer attached to it (**2**). (**3**) Retinal explants were then placed in transwell inserts in a medium/air interphase in which the culture medium was placed under the inserts membrane. (**4**) Samples were maintained for 10 days *in vitro* (10 DIV) changing medium every second day. Treatments (20 µM ML240 or 0.4% DMSO as a vehicle control) were added into the medium on the first day of culture and renewed with every medium change. (**5**) At the end of the culture period, samples were fixed, sectioned, and processed for histological analyses.

When first testing for general photoreceptor cell death markers by TUNEL assay and counting of outer nuclear layer (ONL) cell rows, we did not observe any difference between the different treatment groups (**Figure 2**). This result was expected, since rod photoreceptors in this mouse model are not affected by the mutation, yet represent the majority of photoreceptors in the mouse retina. Only when comparing mutant DMSO-treated explants to untreated WT controls, a higher number of TUNEL-positive cells was observed, which was not found when comparing to the respective mutant control. This can be explained by the overall higher cell death rates in the mutant line.

**Figure 2.**
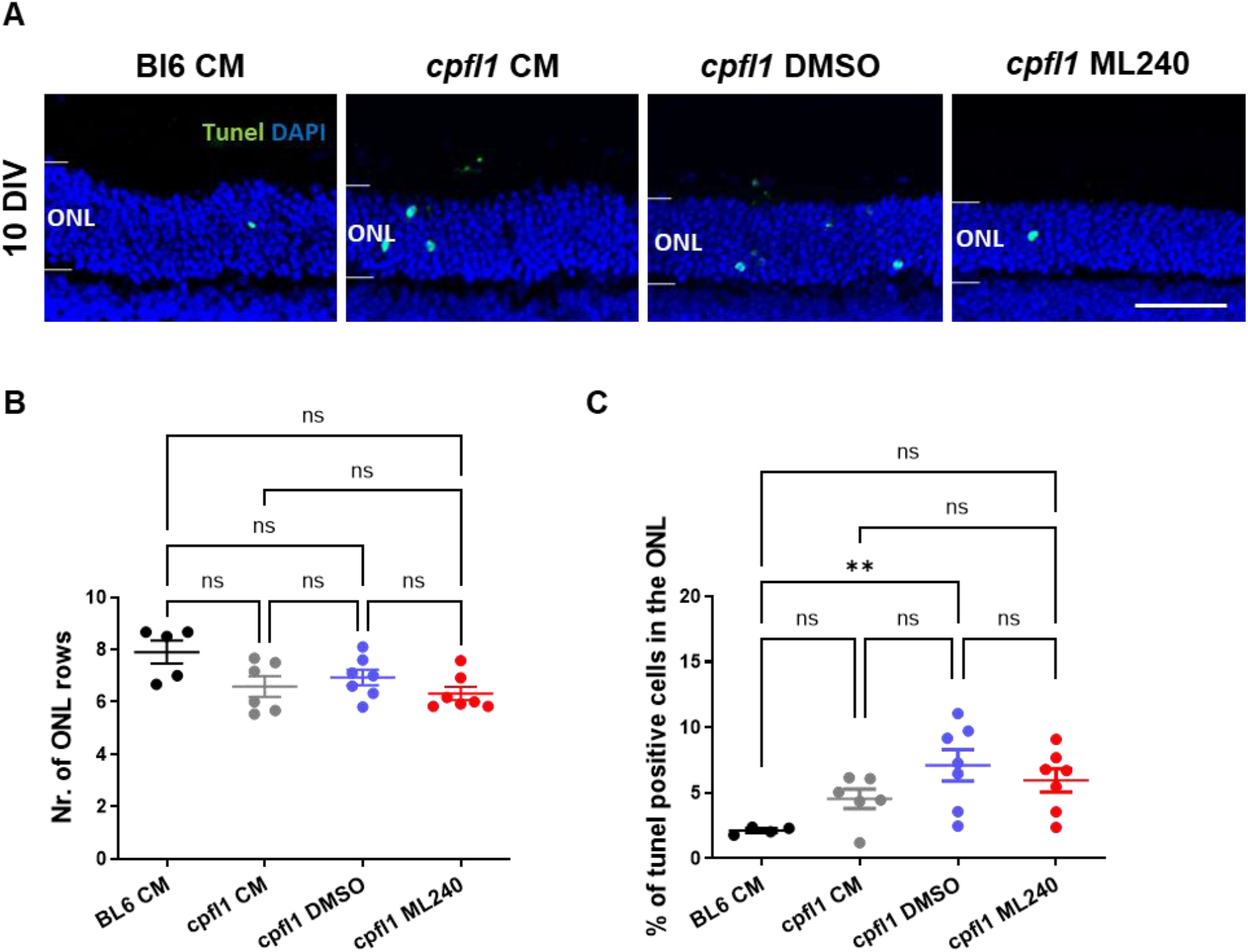
Cell death marker and number of ONL rows remain unchanged in *cpfl1* retinal explants upon VCP inhibition. C57BL/6J or *cpfl1* retinal explants were cultured from PN14 to PN24 (10 DIV). Medium changes and treatments were performed every second day, including the following treatment groups: untreated explants (CM), 0.4% DMSO as a vehicle control, and 20 µM ML240. (**A**) Immunofluorescence images of radial sections of the outer retina visualising TUNEL assay (green) and DAPI (blue) as counterstaining. General photoreceptor cell death was assessed by counting ONL row numbers (**B**) and by quantification of % of TUNEL-positive cells in the ONL (**C)**. Differences between groups were established by one-way ANOVA followed by Tukey’s multiple comparison (n = 4-7). Data presented as mean ± SEM. ns = not significant. ** p < 0.01. Scale bar: 50 µm.

Next, we stained for cone arrestin, a cone photoreceptor-specific marker, to specifically assess the effects of VCP inhibition on cone survival. Reflecting the disease phenotype, *cpfl1* retinal explants showed significantly less cone density than the WT controls, with similar levels in untreated and vehicle-treated controls. However, in *cpfl1* retinal explants treated with 20 µm of the VCP inhibitor ML240, we observed a significantly higher cone density compared with the untreated and vehicle-treated controls (**Figure 3**).

**Figure 3.**
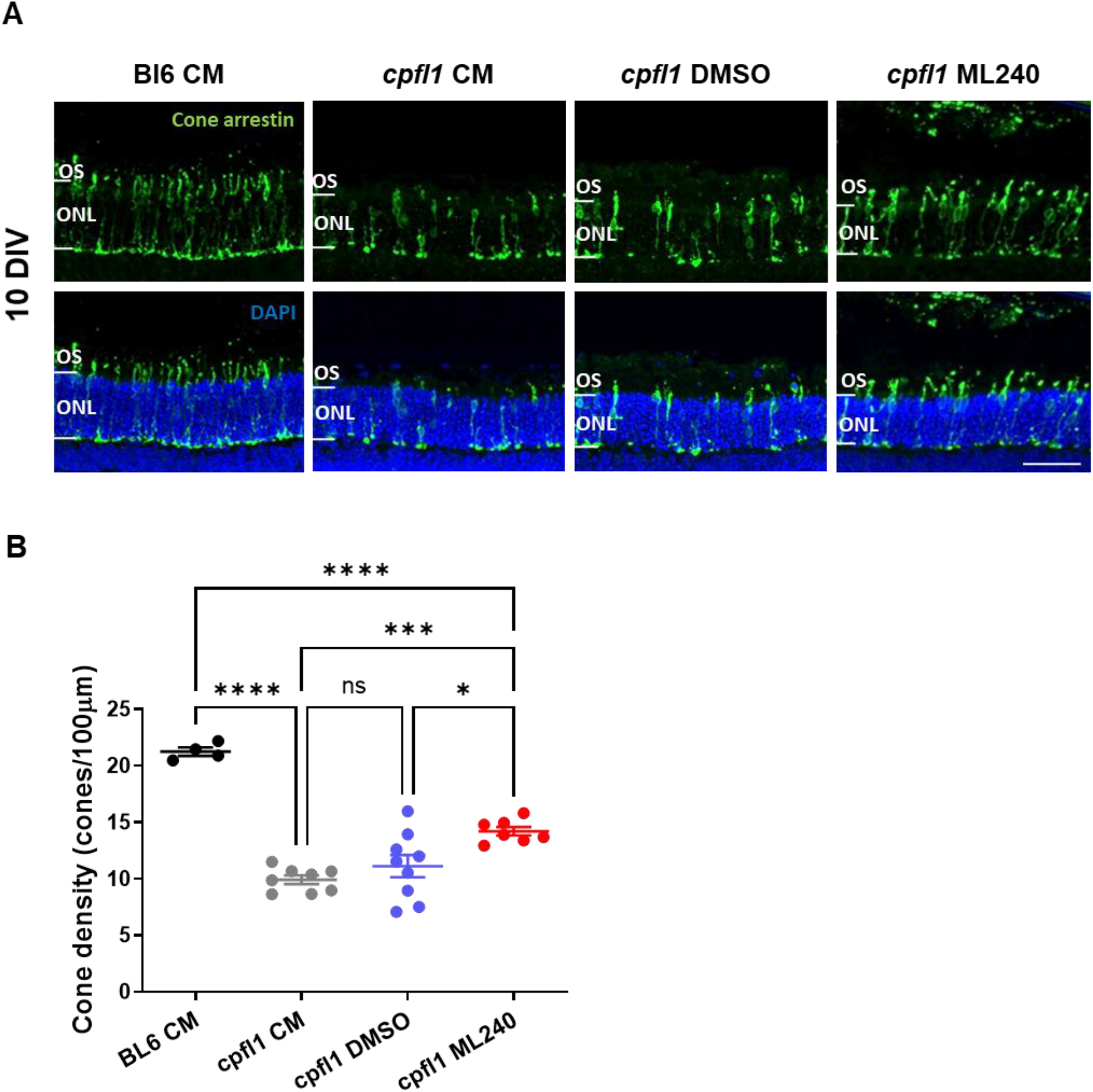
Cone survival is increased after VCP inhibition in *cpfl1* retinal explants. C57BL/6J or *cpfl1* retinal explants were cultured from PN14 to PN24 (10 DIV). Medium changes and treatments were performed every second day, including the following treatment groups: untreated explants (CM), 0.4% DMSO as a vehicle control, and 20 µM ML240. (**A)** Immunofluorescence staining for cone photoreceptors using cone arrestin (green) and DAPI (blue) as counterstaining. Cone survival was assessed as cone density, measured by the number of cones/100 µm of retinal length (**B**). Differences between groups were established by one-way ANOVA followed by Tukey’s multiple comparisons (n = 4-7). Data are presented as mean ± SEM. ns = not significant. * p < 0.05. ** p < 0.01. *** p < 0.001. **** p < 0.0001. Scale bar: 50 µm.

Besides the decrease in the number of cone photoreceptors, a mislocalization of the cone nuclei was reported for the *cpfl1* mouse model. Cone nuclei, in WT retinas normally situated at the most external rows of the ONL, were located towards more internal positions in the inner retina [24]. In our organotypic cultures, this disease phenotype was also reproduced, as we also observed a reduced percentage of cone migration towards the most external part of the ONL in mutant retinal explants compared with the WT. After ML240 treatment, however, this parameter was not significantly improved in the *cpfl1* retinal explants (**Figure 4**).

**Figure 4.**
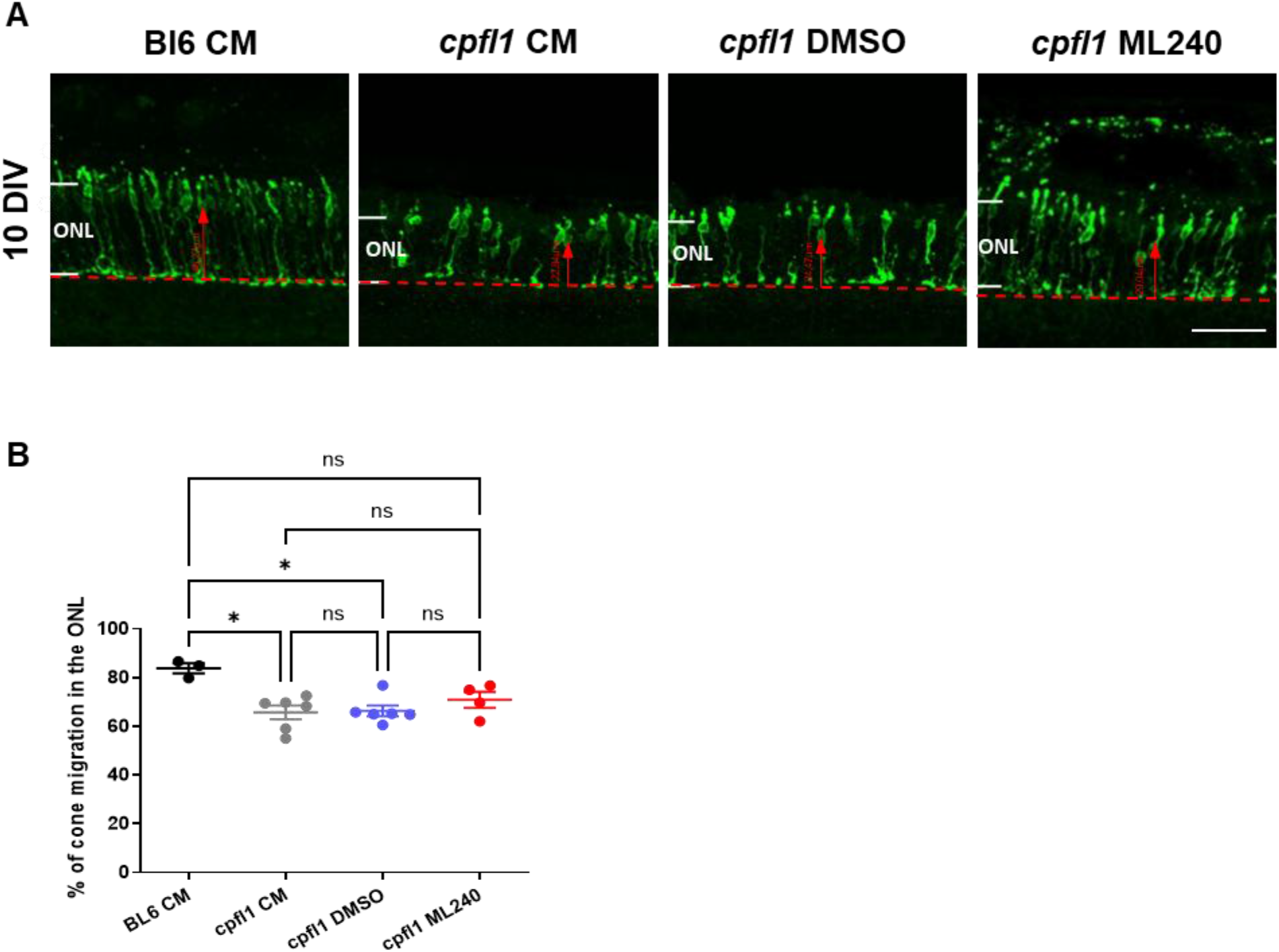
Cone migration through the outer retina is not improved after ML240 treatment. WT or *cpfl1* retinal explants were cultured from PN14 to PN24 (10 DIV). Medium changes and treatments were performed every second day, including the following treatment groups: untreated explants (CM), 0.4% DMSO as vehicle control, and 20 µM ML240. (**A)** Immunofluorescence staining for cone photoreceptors using cone arrestin (green). (**B**) Cone migration through the ONL was assessed by measuring the distance between cone nuclei and the outer plexiform layer (red dotted line), normalised for the thickness of the ONL. Differences between groups were established by one-way ANOVA followed by Tukey’s multiple comparisons (n = 4-7). Data presented as mean ± SEM. ns = not significant. * p < 0.05. Scale bar: 50µm.

### Expression and traffic of cone opsins were improved upon ML240 treatment

After determining that VCP inhibition improved the number of cone photoreceptors in the *cpfl1* model, we further characterized its effects by staining for specific cone opsins, using S and M opsin as markers (**Figure 5**). *cpfl1* CM and DMSO groups showed lower expression of S opsin compared with the WT retinas. Furthermore, we observed an atrophy of cone OSs; fewer cones had outer segments, and in those that did, the segments were shorter. Treatment with ML240 led to a higher expression of S opsin in *cpfl1* explants. While the number of cone photoreceptors with apparent OSs was increased, OSs were also longer compared with the controls, resembling the healthy cone morphology of WT retinal explants. No major differences between the different groups were observed for M opsin expression, which appeared as puncta in some of the cone OSs, often co-localizing with S opsin (**Figure 5**).

**Figure 5.**
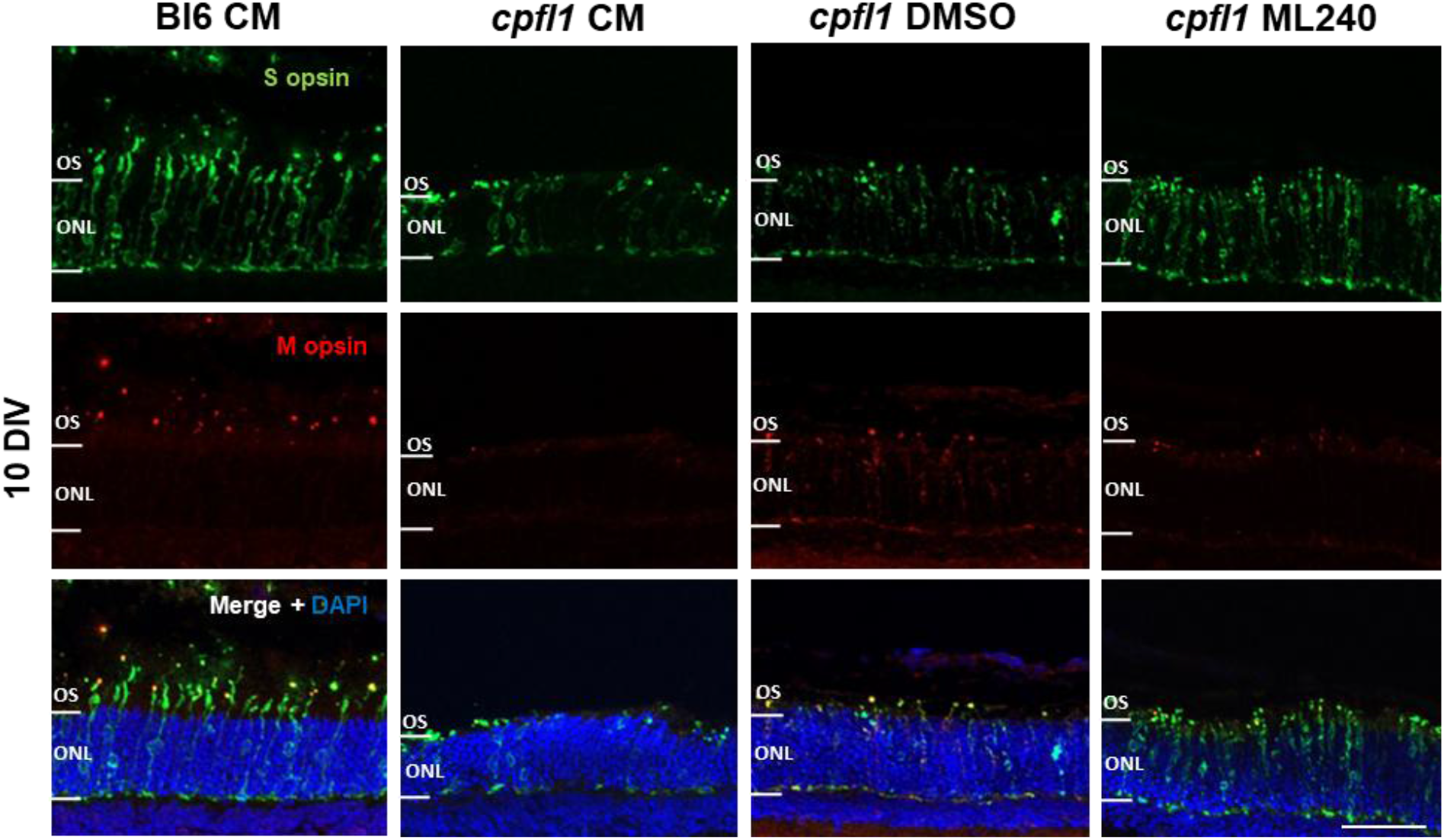
Expression of cone opsins in *cpfl1* retinal explants upon VCP inhibition. C57BL/6J or *cpfl1* retinal explants were cultured from PN14 to PN24 (10 DIV). Medium changes and treatments were performed every second day, including the following treatment groups: untreated explants (CM), 0.4% DMSO as vehicle control, and 20 µM ML240. Immunofluorescence images of S opsin (upper row, green) and M opsin (middle row, red) were obtained. DAPI (lower row, blue) was used for nuclei counterstaining. Scale bar: 50 µm.

### VCP inhibition does not affect the normal visual cycle of healthy rods

As VCP inhibition using ML240 improved cone opsin expression and trafficking to the OSs, we proceeded to check whether the normal trafficking of rhodopsin was altered upon treatment, considering that the visual cycle of rod photoreceptors is not defective in this model. Proper trafficking of rhodopsin to the OSs was not altered in any of the treatment groups compared to the WT condition, indicating that VCP inhibition using ML240 does not disturb the visual function of healthy rod photoreceptors (**Figure 6**).

**Figure 6.**
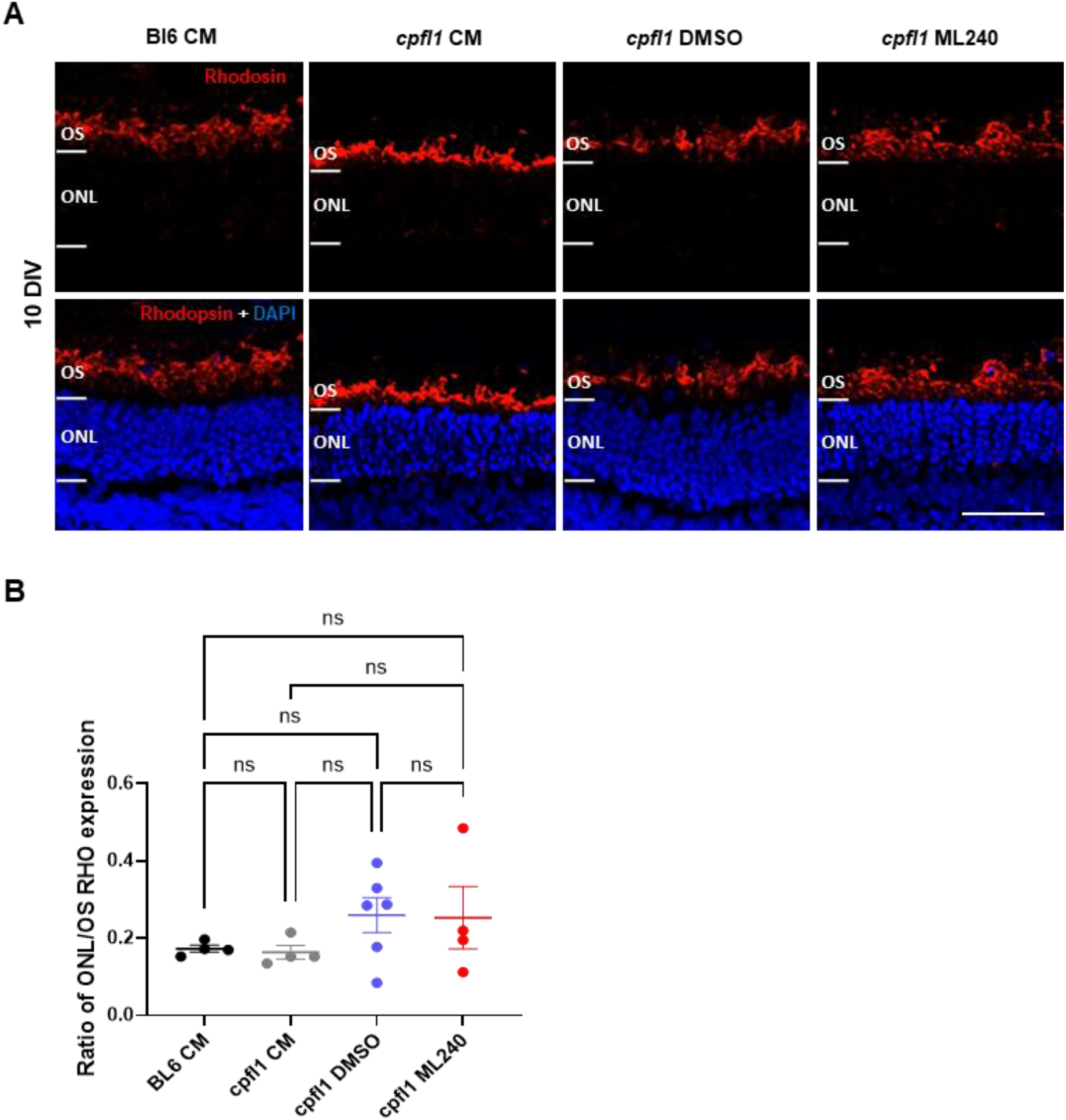
Normal rhodopsin distribution is not affected by ML240 treatment. WT or *cpfl1* retinal explants were cultured from PN14 to PN24 (10 DIV). Medium changes and treatments were performed every second day, including the following treatment groups: untreated explants (CM), 0.4% DMSO as vehicle control, and 20 µM ML240. (**A)** Immunofluorescence staining for rhodopsin (red) and DAPI as nuclei counterstaining. Rhodopsin distribution between the cellular body and outer segments was assessed by the ratio of rhodopsin mean fluorescence intensity between ONL and OS (**B**). Differences between groups were established by one-way ANOVA followed by Tukey’s multiple comparisons (n = 4-7). Data presented as mean ± SEM. ns = not significant. Scale bar: 50 µm.

## Discussion

Achromatopsia (ACHM) is most commonly caused by mutations in genes encoding components of the cone phototransduction cascade, including CNGA3, CNGB3, GNAT2, PDE6H, and PDE6C, as well as ATF6, which is linked to ER homeostasis [3, 25]. Although gene supplementation strategies targeting these individual mutations have shown encouraging results in preclinical models, their success is constrained by the requirement for viable cones at early disease stages and by the mutation-specific nature of these approaches [26].

The present study explores the application of a mutation-independent strategy by targeting a common downstream mechanism of cone degeneration: ER stress and impaired proteostasis [27, 28].

For that purpose, we used the *cpfl1* mouse model, which has a mutation in Pde6c and recapitulates key hallmarks of human ACHM, including elevated cGMP levels and early-onset cone degeneration [6, 20, 24]. Our results showed that pharmacological inhibition of VCP using ML240 significantly preserves cone photoreceptors in an *ex vivo* organotypic culture system. Importantly, this protective effect was specific to degenerating cones, as overall ONL row number and general photoreceptor cell death markers were not significantly altered. This is consistent with the rod-sparing phenotype of the *cpfl1* model, and was expected given that cone photoreceptors represent only∼ 3% of total photoreceptor cells in the mouse retina [20].

Interestingly, besides an improvement in cone density in *cpfl1* retinal explants treated with ML240, we could also observe an improvement in S-opsin expression and outer segment (OS) morphology, known to be disturbed in ACHM models [24]. The improved OS preservation and opsin localization suggest that VCP inhibition not only prevents cell death but also promotes structural maintenance and potentially functional competence of surviving cones.

While cone survival and opsin expression were improved, cone nuclear mislocalization within the ONL was not significantly rescued. This indicates that VCP inhibition may not fully restore developmental or cytoskeletal processes governing cone positioning, or that nuclear displacement represents an earlier or independent pathological event. These findings underscore that structural rescue of cone number and OS integrity does not necessarily equate to complete normalization of retinal architecture. Future studies assessing functional readouts, such as cone-specific electroretinography, will be necessary to determine the extent of functional recovery.

Critical to the translational development of VCP inhibition by ML240 towards clinical applications is safety and the absence of off-target effects. To that end, ML240 treatment did not alter normal rhodopsin trafficking in rod photoreceptors in our model, suggesting that physiological protein transport in healthy rods remains intact under the applied conditions. This observation aligns with our previous reports in other inherited retinal degeneration models, where VCP inhibition reduced ER stress and photoreceptor loss without detrimental effects on unaffected cells [29]. Nonetheless, long-term in vivo studies will be essential to evaluate disease-specific dosing, safety, and potential off-target effects.

Collectively, our results demonstrate that VCP inhibition confers neuroprotection against primary cone degeneration in the *cpfl1* model. By targeting ER stress and proteostasis imbalance, this strategy may overcome limitations associated with gene-specific therapies and provide a broader therapeutic avenue for ACHM and other inherited retinal diseases that may share common downstream pathways leading to photoreceptor cell death. Furthermore, the success of gene therapy approaches depends in part on the presence of intact cone photoreceptors, which can be rescued by gene supplementation [26]. In that regard, the delay of cone photoreceptor cell death observed upon VCP inhibition suggests a potential application of this approach to broaden the therapeutic window in which gene supplementation therapies are effective.

Further in vivo validation and functional analyses will be required to determine the full therapeutic potential of VCP inhibition and its compatibility with emerging gene-based or combination therapies.

